# Varying intolerance of gene pathways to mutational classes explain genetic convergence across neuropsychiatric disorders

**DOI:** 10.1101/054460

**Authors:** Shahar Shohat, Eyal Ben-David, Sagiv Shifman

**Author notes:** Correspondence: Sagiv Shifman, Department of Genetics, The Institute of Life Sciences, The Hebrew University of Jerusalem, Edmond J. Safra campus, Jerusalem 91904, Israel.

## Abstract

Genetic susceptibility to Intellectual disability (ID), autism spectrum disorder (ASD) and schizophrenia (SCZ) often arises from mutations in the same genes, suggesting that they share common mechanisms. We studied genes with *de novo* mutations in the three disorders and genes implicated by SCZ genome-wide association study (GWAS). Using biological annotations and brain gene expression, we show that mutation class explains enrichment patterns more than specific disorder. Genes with loss of function mutations and genes with missense mutations were enriched with different pathways, shared with genes intolerant to mutations. Specific gene expression patterns were found for each disorder. ID genes were preferentially expressed in fetal cortex, ASD genes also in fetal cerebellum and striatum, and genes associated with SCZ were most significantly enriched in adolescent cortex. Our study suggests that convergence across neuropsychiatric disorders stems from vulnerable pathways to genetic variations, but spatiotemporal activity of genes contributes to specific phenotypes.

## Introduction

The successful genetic dissection of Autism spectrum disorder (ASD), intellectual disability (ID) and schizophrenia (SCZ) has resulted in the discovery of a large number of candidate genes (Hamdan et al., 2014; Iossifov et al., 2014; O’Roak et al., 2012; Rauch et al., 2012; De Rubeis et al., 2014). Surprisingly, analysis has suggested that the same genes and the same biological pathways are involved in more than one disorder. First, significant overlap has been found between genes with *de novo* mutations in ID, ASD and SCZ (Fromer et al., 2014; Hoischen et al., 2014; McCarthy et al., 2014). Second, similar pathways have been identified for each of the three disorders: chromatin regulators and synaptic proteins, especially glutamatergic synapses, are pathways affected by genes with *de novo* mutations in ID, ASD and SCZ (Ben-David and Shifman, 2013; Fromer et al., 2014; Hamdan et al., 2014; Hormozdiari et al., 2015; McCarthy et al., 2014; Parikshak et al., 2013; De Rubeis et al., 2014; Willsey et al., 2013). Third, in all three disorders, *de novo* mutations affect genes intolerant to mutations (so-called constrained genes). Specifically *de novo* loss of function (LoF) mutations in ASD and ID are enriched in a set of 1,003 genes that are significantly depleted from mutations in the general human population (and are presumably under the most significant selective constraint) (Samocha et al., 2014).

While analyses of each disorder separately have identified shared pathways between genes, a systematic cross-disorder comparison has not yet been performed. Such an analysis could yield further insight into the shared, convergent etiology, but critically it could also identify distinct, divergent features that are hallmarks of genes associated with each of the different disorders.

In this study, we investigate the convergence and divergence between genes associated with ID, ASD and SCZ using a methodology applied for individual disorders (Ben-David and Shifman, 2012; Gilman et al., 2011, 2012; Gulsuner et al., 2013; O’Roak et al., 2012; Willsey et al., 2013). To make the analyses comparable and to avoid ascertainment bias, we studied genes identified using the same experimental strategy in each disorder. We focused on functional *de novo* mutations from exome sequencing studies as they represent a set of prime candidate genes. Our analyses revealed specific pathways that are associated with LoF mutations and different pathways with missense mutations across disorders. Despite the shared attributes, unique patterns of expression in the brain were identified for genes disrupted by *de novo* mutations in ASD and ID, as well as for genes in GWAS loci for SCZ.

## Results

### Enrichment of de novo mutations depending on diagnosis, mutation class and sex

We collected sets of genes with coding *de novo* SNVs identified by genome wide screens in ID, ASD and SCZ (Table S1). The data included exome sequencing of 195 ID, 3,953 ASD, and 1,027 SCZ cases. As a control, we included *de novo* mutations identified in unaffected siblings of ASD or SCZ individuals (n = 1,995), and unrelated typically developing individuals (n = 34). These mutations were divided into three classes based on their expected severity: (1) LoF (nonsense, splice site and frameshift), (2) missense (non-synonymous SNV and indels that did not result in a frameshift), and (3) synonymous mutations. Since most synonymous mutations are expected to be silent, across this study we treated the synonymous mutations as a negative control for the mutations more likely to be functional (LoF and missense). Our strategy, focusing on *de novo* mutations, excludes recessive mutations, which were mostly identified for ID by homozygosity mapping (Najmabadi et al., 2011).

We calculated for each disorder and each mutation class the mutation rate per individual and tested the ratios between functional and non-functional mutation rates to correct for experimental confounders. We replicated the higher rate of *de novo* mutations in ID and ASD, and found a significant enrichment for missense mutations in SCZ that was not reported before (Figures 1A and S1A). LoF mutations were significantly enriched in ID (FDR corrected *P* = 5.5×10^−8^) and ASD (FDR corrected *P* = 6.0×10^−7^), but not in SCZ (FDR corrected *P* = 0.063). For missense mutations, the ratio was significantly higher compared to the control in ID (FDR corrected *P* = 4.7×10^−5^), ASD (FDR corrected *P* = 3.1×10^−4^), and SCZ (FDR corrected *P* = 2.3×10^−4^). Consistent with a previous study (Fromer et al., 2014), the rates of both LoF and missense mutations were higher in ID compared to ASD (FDR corrected *P_LoF_* = 1.7 × 10^−6^, FDR corrected *P_missense_* = 3.1 × 10^−4^) and SCZ (FDR corrected *P_LoF_* = 6.0 × 10^−7^, FDR corrected *P_missense_* = 0.0027). No significant difference was found between ASD and SCZ (FDR corrected *P_LoF_* = 0.17, FDR corrected *P_missense_* = 0.090).

**Figure 1.**
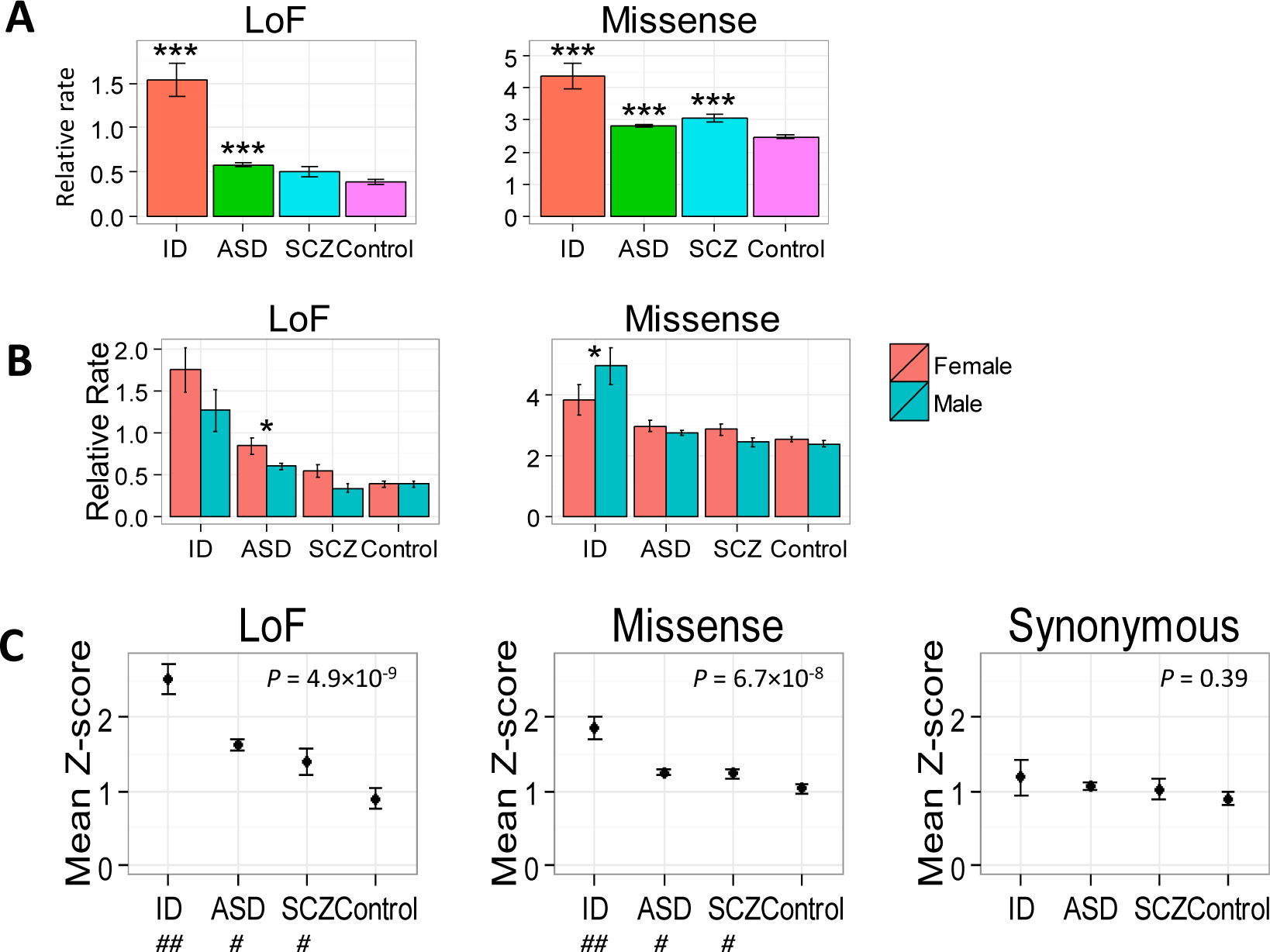
Rates of *de novo* mutations and levels of constraint for genes with *de novo* mutations (See also Figure S1). (A) The per-individual rate of functional mutations (LoF or missense) relative to the study-specific rate of non-functional mutations (synonymous). Error bars are standard error of the mean (SEM). *P*-values represent the significance of the difference from the controls. *P*-values were corrected for multiple testing with the Benjamini and Hochberg false discovery rate (FDR) procedure. ***, FDR corrected *P* < 0.001. (B) Differences between females and males in the rate of *de novo* mutations (values as above). *P*-values represent significant interaction between sex and disease status. *, FDR corrected P < 0.05. (C) The mean level of constraint (Z-score) ± SEM for genes with different types of mutations in the three disorders and the control. *P*-values were obtained by one-way ANOVA test for equal means. Post-hoc test can be found in table S3. #, significantly different from control. ##, significantly different form all other disorders.

Previous studies in ASD reported a higher rate of *de novo* SNVs in females compared to males (Iossifov et al., 2014; De Rubeis et al., 2014). We found evidence that *de novo* LoF mutations were at higher rate in females not only in ASD, but also across disorders (all uncorrected *P* < 0.05; FDR corrected *P* < 0.05 only for ASD). Surprisingly, *de novo* missense mutations were at significantly higher rate in males with ID, but not in ASD or SCZ (Figures 1B and S1B).

We next tested the overlap of genes between disorders. We found a significant overlap for both LoF (observed to expected ratio [O/E] = 2.94-15.24) and missense mutations (O/E = 2.24 – 2.80) between disorders (Table S2). The most significant overlap for genes with LoF mutations was between ID and ASD (O/E = 15.24), and for genes with missense mutations between SCZ and ASD (O/E = 2.80). In some pairwise comparisons, there was a small but significant overlap between genes identified in the disorders and the controls (O/E = 0-1.8). For synonymous mutations, there was no significant overlap between conditions (all *P values* > 0.05).

### Genes with missense mutations have significantly more protein–protein interactions while genes with LoF mutations show less variation in gene expression

There are differences between genes in the sensitivity to functional mutations. We compared the degree of intolerance to mutation (constraint) for genes with different class of *de novo* mutations and between the disorders. For genes with LoF and missense mutations there was a significant difference in the average constraint score among the different conditions (*P_LoF_* = 4.9×10^−9^, *P_missense_* = 6.7×10^−8^), but this was not significant for synonymous mutations (*P* = 0.39) (Figure 1C). The average constraint score for genes with LoF and missense mutations was significantly higher for ID, following by ASD and SCZ, which were not significantly different from each other, but were significantly different from the controls (Post-hoc tests, Table S3). The overlap between previously defined constrained genes (n = 1003) (Samocha et al., 2014) and the genes with mutations was significant across the three disorders for LoF and missense mutations (all FDR corrected *P* < 0.05), but not for genes with synonymous mutations (all FDR corrected *P* > 0.05) (Figure S1C).

Since genes mutated in ID, ASD and SCZ are under higher selective constraint, we asked why those genes were sensitive to mutations. We hypothesized that the genes may code for proteins having more protein–protein interactions (PPIs). It has been previously shown that mutations in highly connected proteins (hub proteins) are more likely to be lethal (Jeong et al., 2001). Consistent with our hypothesis, we found a significant positive correlation between the median weighted number of interactions and the level of constraint across disorders and mutation types (Pearson correlation r = 0.92, *P* = 2.4×10^−5^, Figure 2A). The level of constraint and the number of interactions were also positively correlated across all genes (r = 0.24, *P* < 2.2×10^−16^), and the number of interactions for constrained genes was significantly higher compared to non-constrained genes (*P* < 2 × 10^−16^) (Figure 2B).

**Figure 2.**
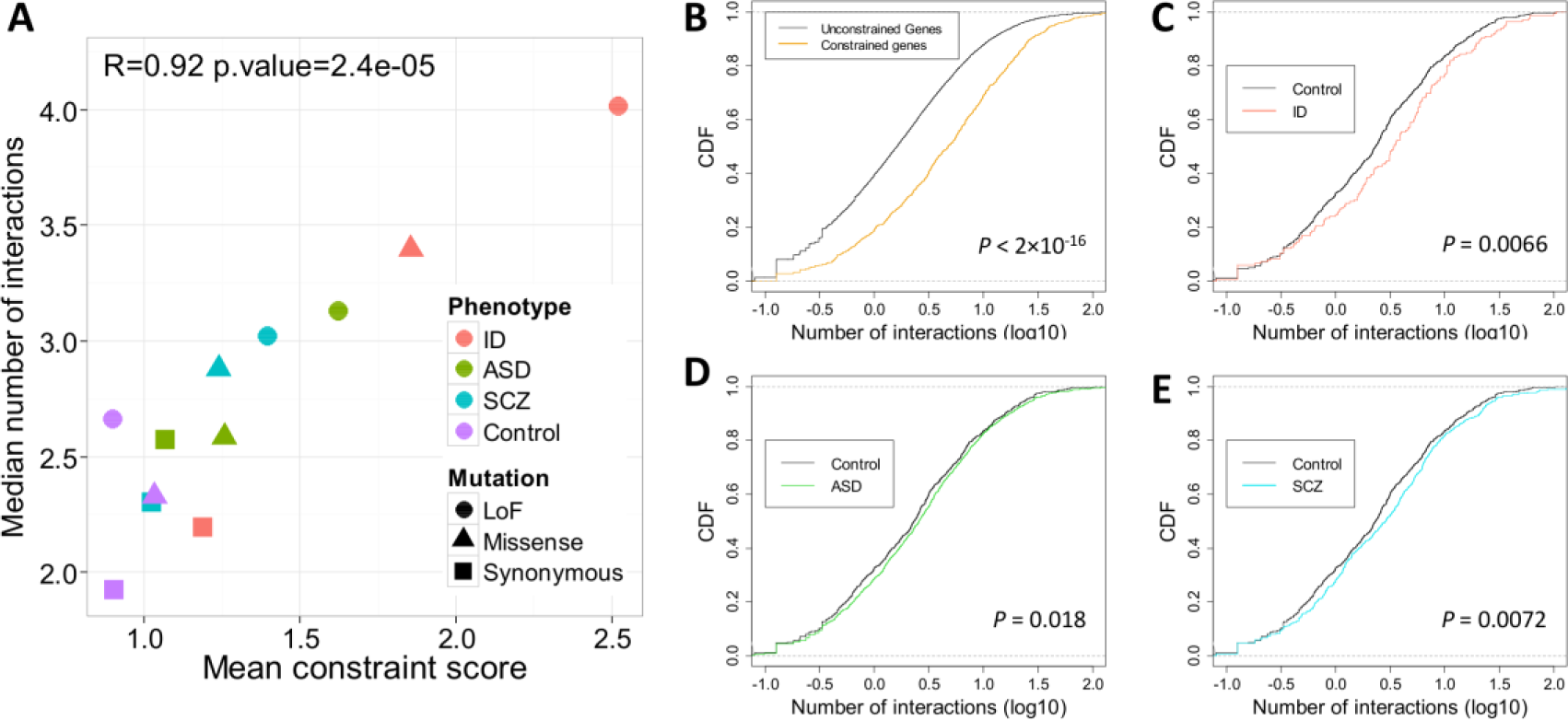
Constrained genes and genes with functional *de novo* mutations in the disorders tend to have more protein-protein interactions (See also Figure S2). (A) Positive correlation (r = 0.92) between the median number of interactions and the mean constraint score across different mutation categories and disorders. (B) Cumulative distribution function (CDF) of the number of interactions (log_10_) plotted for constrained and unconstrained genes. The *P*-values were calculated using a two-sample Kolmogorov–Smirnov test. (C-E) CDF of the number of interactions (log_10_) plotted for genes with missense mutations in the control relative to ID, ASD or SCZ. *P*-values were calculated using a two-sample Kolmogorov–Smirnov test, and corrected by FDR.

We expected the genes with functional mutations to be enriched with hub proteins, but we found this to be significant only for missense mutations. We tested the difference between the number of interactions per protein in the three disorders compared to the control. For genes with synonymous and LoF mutations, there was no statistical difference between any of the three disorders and the control (Figure S2). In contrast, there was significant higher number of interaction partners for genes with missense mutations in ID, ASD and SCZ (FDR corrected *P_ID_* = 0.0066, *P_ASD_* = 0.018, *P_SCZ_* = 0.0072) (Figure 2C-E).

Pathogenic missense mutations may be especially associated with hub proteins, because changes in the protein sequence may lead to changes in the ability to interact with other protein partners. Genes may also be under strong selective constraint if the level of expression is critical for proper function (dosage-sensitive genes). Under this hypothesis, we expect constrained genes, as well as genes with pathogenic LoF mutations to show low variation in gene expression (Hasegawa et al., 2015). To test this, we studied previously published single-cell gene expression from human brain (Darmanis et al., 2015). For each gene, we calculated a standardized measure of expression variation across single-cell neurons (based on coefficient of variation (CV) controlled for mean expression). Consistent with our prediction, constrained genes had on average a significantly lower variation in expression relative to unconstrained genes (*P* < 2 × 10^−16^) (Figure 3). Similarly, we observed across the disorders a trend of lower variation in gene expression for genes mutated by functional mutations, with a stronger effect size for LoF mutations (Figure 3). When testing all the disorders together, genes with LoF mutations have a significantly lower expression variation relative to genes with missense mutations (Figure S3). Thus, our analysis supports a pathogenic role in all three disorders for mutations in genes intolerant to functional mutations, possibly because some of those genes code for ‘hub’ proteins or proteins sensitive to changes in expression levels (haploinsufficient).

**Figure 3.**
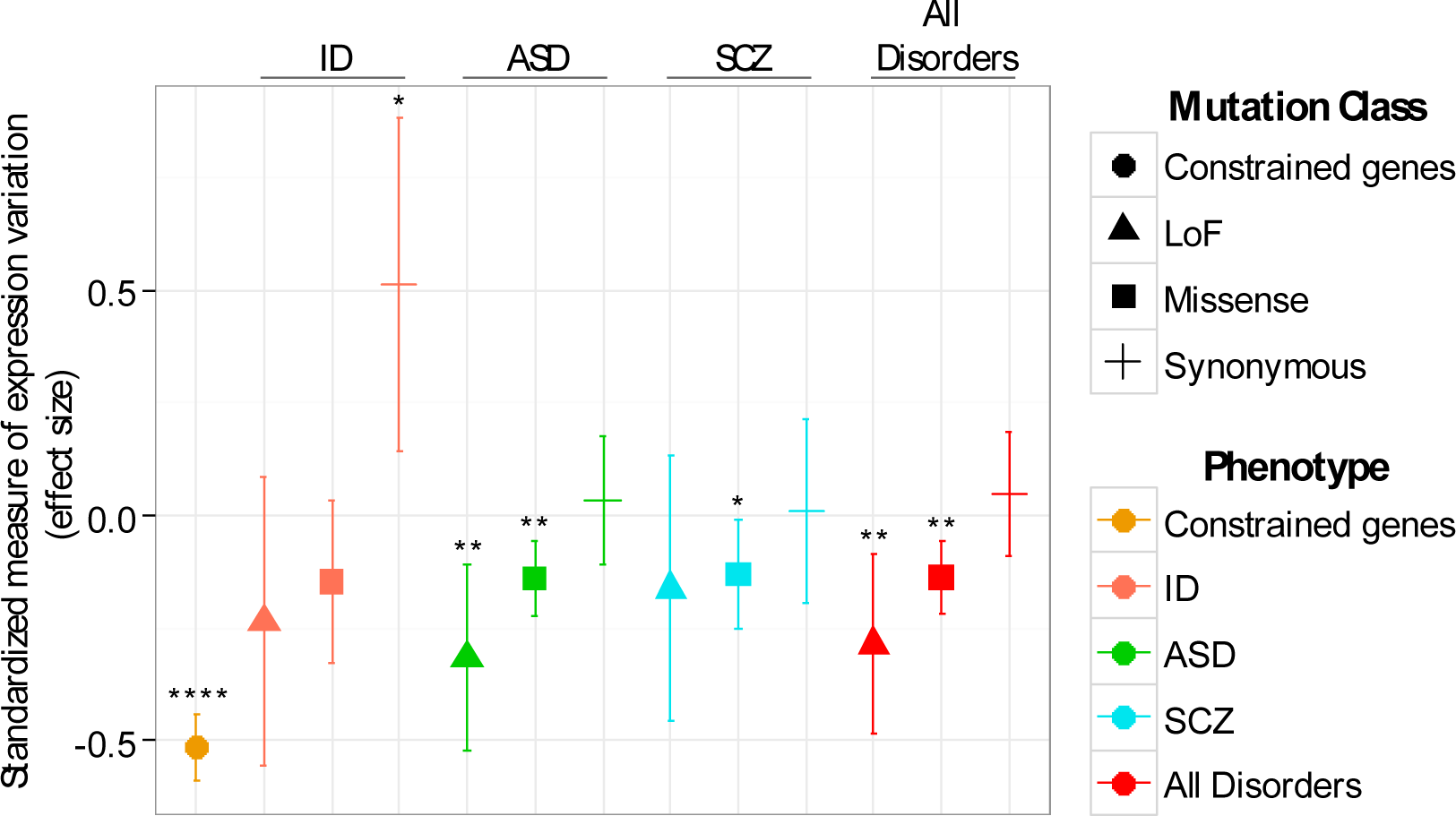
Constrained genes and genes with functional mutations tend to have lower variation in gene expression (See also Figure S3). Values are the effect size (Cohen’s d) for the difference in the standardized measure of expression variation between genes with mutations in the disorders and the control (± 95% confidence interval). *, FDR corrected P < 0.05. **, FDR corrected P < 0.01. ****, FDR corrected P < 0.0001.

### Constrained genes are involved in generation of neurons, expressed in multiple brain regions, but most significantly in early mid-fetal cortex

Since constrained genes are not linked to any specific phenotype, we wanted to characterize their biological processes and expression patterns in the brain. We found that the constrained genes were enriched for biological processes that included generation of neurons, axon development, neurogenesis, and chromatin modification (FDR corrected *P* < 5 × 10^−31^) (Figure 4A). In addition, the constrained genes were significantly associated with neurodevelopmental phenotypes in humans (based on the Human Phenotype Ontology), including neurodevelopmental abnormality, global developmental delay and cognitive impairment (FDR corrected *P* < 6 × 10^−11^).

**Figure 4.**
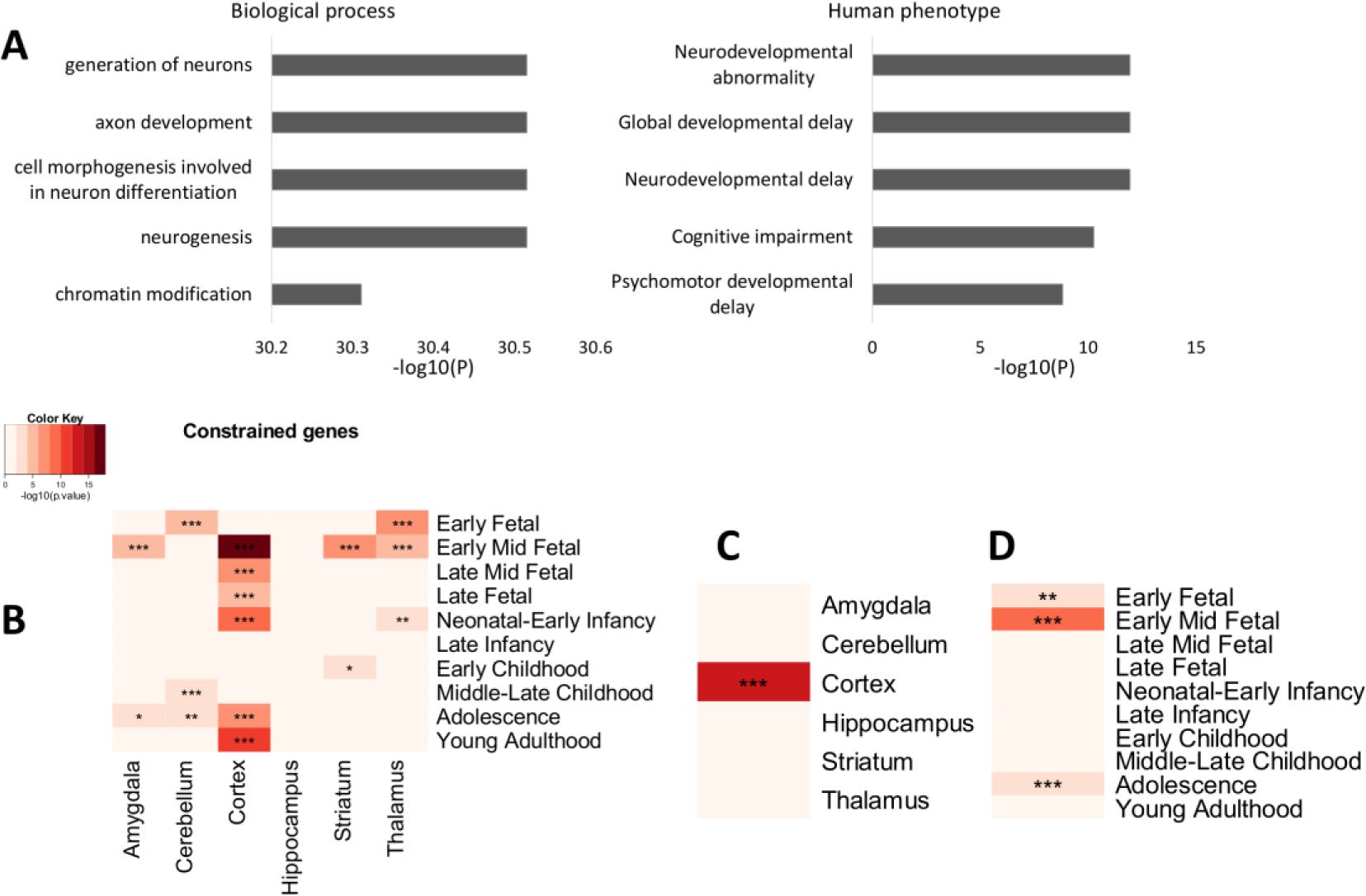
Constrained genes are involved in specific biological processes and show specific spatiotemporal patterns of expression in the brain. (A) Biological processes (left) and human phenotypes (right) that are significantly enriched in constrained genes. The analysis was based on the pathway enrichment tool ToppGene Suite, which includes phenotypes from the Human Phenotype Ontology (HPO). Values are ‐ log_10_ of the FDR corrected *P*-values. (B-D) Enrichment in brain regions and during specific developmental epochs. Darker colors signify higher enrichment, presented as −log_10_ of the *P*-values. *P*-values are calculated using the Fisher’s exact test, and are corrected for multiple tests using FDR. *, FDR corrected P < 0.05. **, FDR corrected P < 0.01. ***, FDR corrected P < 0.001. The enrichment analysis was performed on (B) expression in brain regions during different developmental stages, (C) expression in different brain regions, averaged across development stages, and (D) expression in different developmental stage averaged across brain regions.

We next evaluated whether constrained genes are enriched in specific brain regions and developmental stages. We used a previously published method (Dougherty et al., 2010; Xu et al., 2014) that gives a measure of enrichment of a gene in a tissue or cell type (specificity index probability (pSI) statistic). We used a pSI dataset based on human brain gene expression to test for enrichment of constrained genes in six different brain region (amygdala, cerebellum, cortex, hippocampus, striatum, thalamus) and ten developmental periods (from early fetal to young adulthood). Consistent with previous study (Wells et al., 2015), the constrained genes were significantly enriched (FDR corrected *P* < 0.05) across many brain regions (all regions excluding the hippocampus) and across different stages of development, but mid-fetal cortex was the most significant (FDR corrected *P* = 2.6 × 10^−17^) (Figure 4B). The cortex stood up as the brain region most enriched with constrained genes (FDR corrected *P* = 1.1 × 10^−13^) (Figure 4C), as well as three critical developmental periods, early fetal, early mid-fetal and adolescence (FDR corrected *P* = 2 × 10^−8^-4 × 10^−3^) (Figure 4D).

### Preferential expression of genes mutated in ID and ASD in specific brain regions and developmental stages

We next studied the enrichment of genes mutated in the disorders across multiple brain regions and developmental stages (Figure 5). In the case of ID, the enrichment was restricted to the cortex. ID genes with LoF mutations were most significantly enriched in early mid-fetal cortex (FDR corrected *P* = 0.024) (Figure 5A), while genes with missense mutations showed the strongest enrichment in young adulthood cortex (FDR corrected *P* = 0.0060) (Figure 5B). Genes with missense mutations in ASD did not show significant enrichment in any brain region. Consistent with previous studies (Parikshak et al., 2013; Willsey et al., 2013), genes with LoF mutations in ASD showed a significant enrichment in early mid-fetal cortex (FDR corrected *P* = 0.0019) (Figure 5C), a region that was the most significantly associated with ID genes disrupted by LoF mutations and constrained genes.

**Figure 5.**
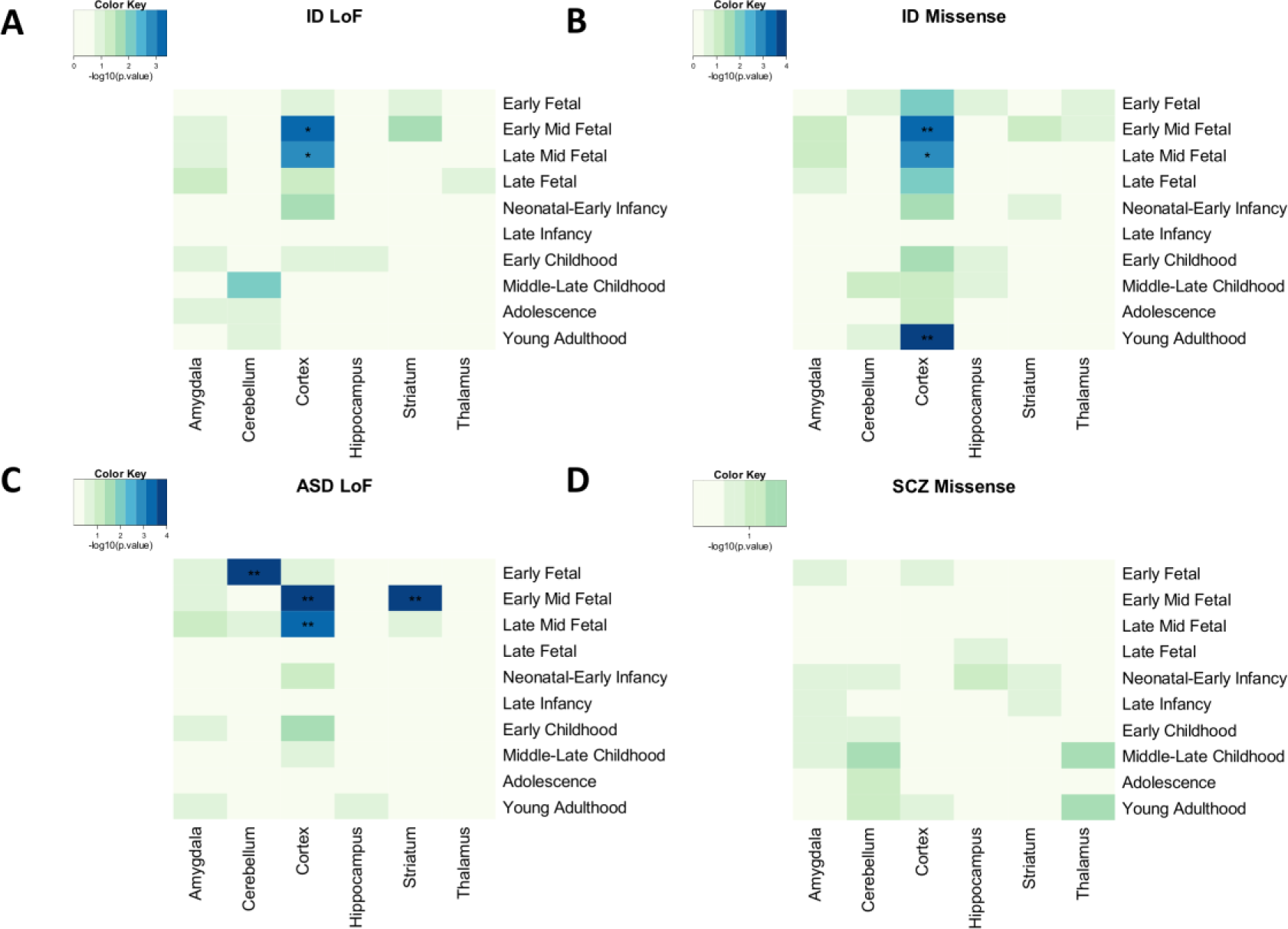
Region and stage specific enrichment in the human brain for genes with functional *de novo* mutations in ID and ASD (see also Figure S4). Heatmap colors represent the enrichment significance presented as −log_10_ of the *P*-values. *P*-values are calculated based on permutations in the *dnenrich* software and are corrected for multiple tests using FDR. *, FDR corrected P < 0.05. **, FDR corrected P < 0.01. (A) Enrichment analysis of genes with LoF mutations in ID. (B) Enrichment analysis of genes with missense mutations in ID. (C) Enrichment analysis of genes with LoF mutations in ASD. (D) Enrichment analysis of genes with missense mutations in SCZ.

However, in contrast to ID, the enrichment in ASD was not restricted to the cortex. Two additional regions were enriched in ASD genes with LoF mutations: early mid-fetal striatum (FDR corrected *P* = 0.0019) and early fetal cerebellum (FDR corrected *P* = 0.0019) (Figure 5C). Since there is a significant overlap between genes with LoF mutations in ASD and ID, we retested the enrichment excluding the overlapping genes. Despite the smaller number of non-overlapping genes in ID and ASD with LoF mutations (n_ID_ = 39, n_ASD_ = 506), the enrichment pattern did not change in a noteworthy way in both cases (Figure S4). Unlike ASD or ID, in SCZ none of the regions showed significant enrichment, after correcting for multiple testing. The most nominally significant enrichment was in the cerebellum during middle-late childhood, when testing genes with missense mutations (nominal *P* = 0.030, FDR corrected *P* = 0.62) (Figure 5D). Gene with synonymous mutations and genes with mutations in the control did not show any significant enrichment.

We next analyzed the genes mutated in the three disorders for enrichment of biological processes (Figures S4C and S4D). Genes with LoF mutations in ID were enriched for chromosome organization and regulation of transcription (FDR corrected *P* = 0.013). Genes with LoF mutations in ASD were enriched for similar biological processes, including chromatin modification (FDR corrected *P* = 1.6×10^−10^) and chromosome organization (FDR corrected *P* = 4.7×10^−8^). Genes with LoF mutations in SCZ did not show any significant enrichment. Genes with missense mutations across disorders were significantly enriched with similar and overlapping biological processes related to neuron development (FDR corrected *P* = 3.4×10^−5^ − 1.7×10^−13^) (Figure S4D).

### The analysis of gene co-expression network in the developing human cortex show similar patterns for ID and ASD genes, strongly depending on mutation class

Our analysis points to enrichment in fetal cortex for genes with functional mutations in both ID and ASD. Since previous studies showed that ASD genes are co-expressed in fetal cortex, we wanted to test the specificity of the enrichment by comparing between ID and ASD and between different mutation classes. We used a weighted gene co-expression network analysis based on gene expression from cortical development that divided the genes into 18 different modules (labeled M1 ‐ M18) (Parikshak et al., 2013). The statistical power to detect enrichment may differ considerably between disorders as the power is a function of the degree of enrichment, but is also a function of the sample size (number of genes in each category) and the fraction of genuine risk genes among the list of mutated genes. To alleviate this problem and directly assess the convergence of mutations in the different disorders to the same modules, we calculated the correlation between the enrichment strength in each module across the different disorders and the different mutation categories. We used the correlation as the input for hierarchical clustering, allowing us to globally survey how related are the different disorders and mutation classes (Figure 6A).

**Figure 6.**
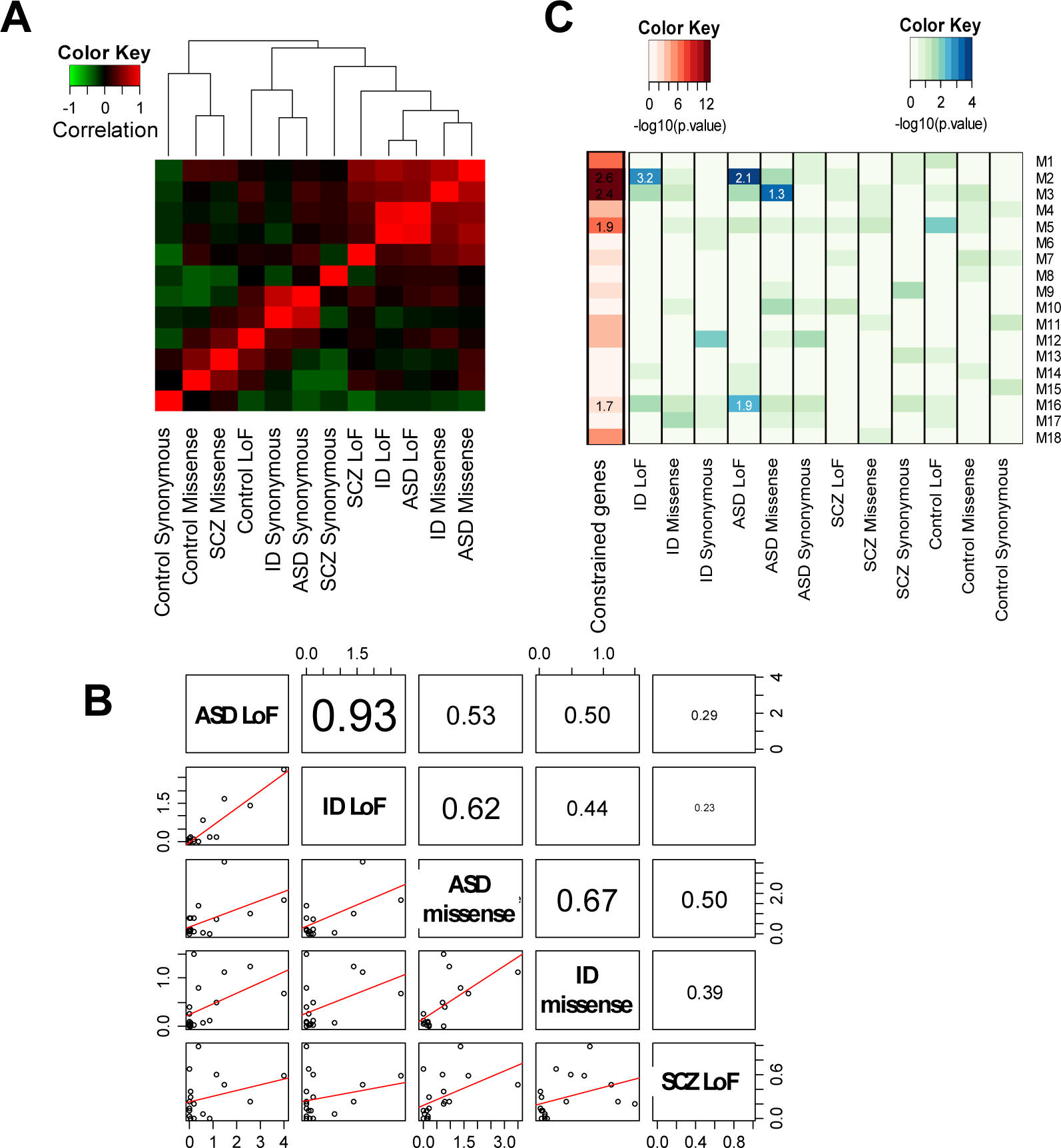
Coexpression in the developing cortex of genes mutated by *de novo* mutations in ASD, SCZ, ID and unaffected control (see also Figure S5). (A) The correlations between disorders and mutation classes are visualized as a heatmap. Samples are sorted using hierarchical clustering. Red colors signify positive correlation and green negative correlation. (B) A scatter plot showing the correlation between mutations classes and disorders. Below the diagonal are plots of the *P*-values (-log_10_) for each module in one condition & mutation class as a function of another. The red line is the linear best fit. Above the diagonal are the correlation values, with the size of the font proportional to the correlations strength. (C) Enrichment across 18 WGCNA modules of genes with mutations. The enrichment is shown for constrained genes and for different disorders divided to different classes of mutations. Statistical significance of enrichment of *de novo* mutations within each module is calculated with the *dnenrich* software (Fromer et al., 2014). Heatmap colors represent −log_10_ of the nominal *P*-values. The numbers are enrichment odds ratios, shown only for significant (FDR corrected) and positive enrichments.

We observed a strong relationship between ASD and ID genes highly influenced by the mutation class (Figure 6A). Specifically, the most significant correlation is observed between genes with LoF mutations in ID and ASD (r = 0.93, FDR corrected *P* = 1.5×10^−7^) (Figure 6B). This correlation across disorders (r = 0.93) was significantly higher than the correlations within disorders between LoF and missense mutations (*P*_ASD_ = 0.0018, *P*_ID_ = 0.0006); correlations which were non-significantly different than 0 (FDR corrected *P* > 0.05, r_ASD_ = 0.53, r_ID_ = 0.44). Similarly, genes with missense mutations in ASD are most correlated with missense mutation in ID (r = 0.67, FDR corrected *P* = 0.015).

When looking at specific modules (Figure 6C), the most significant enrichment was in the M2 module for genes with LoF mutations in ID (FDR corrected *P* = 0.029) and ASD (FDR corrected *P* = 0.0018), as well as for constrained genes (FDR corrected *P* = 3.2×10^−12^). Based on functional annotations the M2 module is enriched for chromatin modification (FDR corrected *P* = 8.4×10^−12^), chromosome organization (FDR corrected *P* = 8.8×10^−9^), and genes known to cause embryonic lethality in mice (FDR corrected *P* = 4.2×10^−6^). Across all modules, there was no significant enrichment for SCZ or for genes with synonymous mutations, nor for mutations in the control (Figure 6C).

A previous study included genes with recessive inheritance in the analysis of ID (Parikshak et al., 2013). Based on the sensitivity of our analysis to mutation class, our hypothesis was that genes sensitive to mutations in both copies would be involved in different biological processes. To explore this possibility we tested the enrichment of ID autosomal recessive genes in the WGCNA modules, as well as for functional annotations. Consistent with our hypothesis, the recessive ID genes were not significantly enriched in any module (all FDR corrected *P* > 0.05), and were enriched for very different functional annotations. The biological processes annotations enriched for autosomal recessive ID genes were mainly related to metabolism (e.g. organic acid metabolic process and oxoacid metabolic process) (Figure S5). A similar functional annotation enrichment for metabolic process was observed for genes associated with autosomal recessive inheritance annotation in the Human Phenotype Ontology (HPO).

### Genes implicated by schizophrenia genome-wide association preferentially expressed in the cortex during adolescence and in DRD2 medium spiny neurons

Since the analysis of *de novo* mutations in SCZ did not yield a conclusive enrichment in the brain, we turned to analyze common variants associated with SCZ. A genome-wide association study (GWAS) of 36,989 SCZ cases identified 108 loci significantly associated with a small increase in SCZ risk (Ripke et al., 2014). Many of the loci identified by SCZ GWAS contain multiple SNPs in linkage disequilibrium and more than one gene. We therefore used a multistep procedure to prioritize the most likely causal genes.

Testing the list of genes for enrichment of functional annotations revealed a significant enrichment of multiple relevant biological processes and cellular components, such as synaptic transmission, generation of neurons, somatodendritic compartment, and synapse (FDR corrected *P* < 8 × 10^−4^) (Figure S6). The 60 genes that were included in at least one of the significant annotations were considered as the most likely SCZ candidate genes (Table S4). We found that the 60 genes are preferentially expressed in specific brain regions, developmental stages, and in neural cell types. Across brain regions and developmental stages the most significant enrichment was for the cortex during adolescence and young adulthood (FDR corrected *P* = 0.004) (Figure 7A), and across cells for DRD2+ (FDR corrected *P* = 0.008) and DRD1+ medium spiny neurons (FDR corrected *P*= 0.01) (Figure 7B).

**Figure 7.**
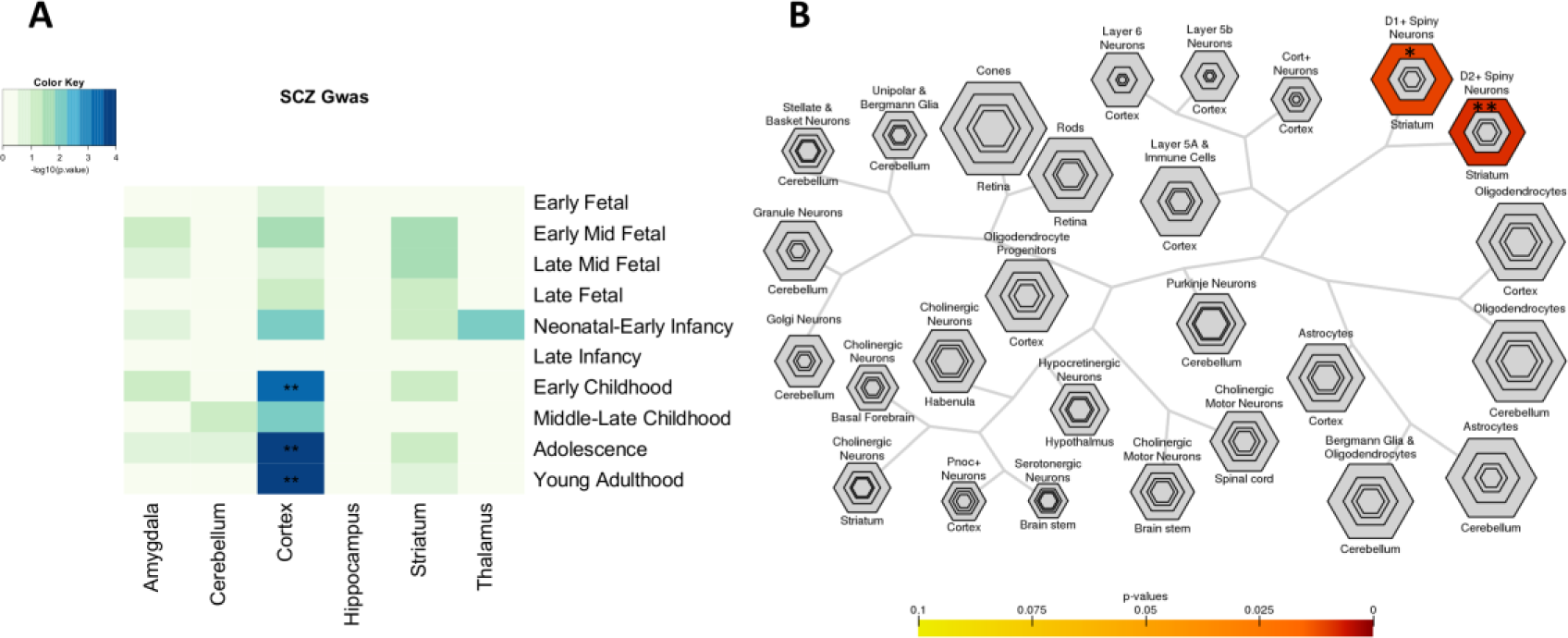
Preferential expression in the human brain for neuronal genes associated with SCZ (see also Figure S6). Heatmap colors represent the enrichment significance presented as −log_10_ of the *P*-values. *P*-values are calculated based on Fisher’s exact test corrected for multiple tests using FDR. *, FDR corrected P < 0.05. **, FDR corrected P < 0.01. (A) Analysis of adult brain regions and development. (B) Analysis across cell types is shown for different specificity Index thresholds (pSI). The outer hexagons represent a pSI < 0.05 and the inner hexagons a more stringent pSI. The size of the hexagons is scaled to the size of the gene list.

## Discussion

Our analyses of genes that contain *de novo* mutations in ID, ASD and SCZ revealed the existence of different enrichment patterns for different classes of mutations. This enrichment is not related to a specific disorder. Table 1 summarizes our findings and shows the molecular and neuronal processes most vulnerable to specific types of mutation. The three disorders show divergence in the specific brain regions and developmental periods affected by genetic variations, consistent with differences in their age of onset.

**Table 1.** Features enriched in genes with specific types of genetic variants

|  |  | ID | ASD | SCZ | Constrained |
| --- | --- | --- | --- | --- | --- |
| **LoF** | Mutation rate relative to control | *** | *** | NS | - |
|  | GO: chromatin/chromosome organization | * | *** | NS | *** |
|  | Low variance in expression | NS | ** | NS | *** |
|  | Expression: Mid-fetal cortex | * | ** | NS | *** |
|  | WGCNA: M2 module | *** | *** | NS | *** |
| **Missense** | Mutation rate relative to control | *** | *** | *** | - |
|  | GO: neuron development | *** | *** | *** | *** |
|  | Hub proteins | ** | * | ** | *** |
|  | Expression: Young adulthood cortex | ** | NS | NS | *** |
|  | WGCNA: M3 module | NS | ** | NS | *** |
| **GWAS** | GO: synaptic transmission | NA | NA | *** | *** |
|  | Expression: Adolescence cortex | NA | NA | ** | *** |

\*, FDR corrected $P < 0.05$ . \*\*, FDR corrected $P < 0.01$ . \*\*\*, FDR corrected $P < 0.001$ . NS, non-significant. NA, data not available. GO, gene ontology. Expression, enrichment based on gene expression in the brain. WGCNA, weighted gene coexpression network analysis. Hub proteins, proteins with a large number of interactions. ID, intellectual disability. ASD, autism spectrum disorder. SCZ, schizophrenia. Constrained, constrained genes.

Our results point to molecular convergence across disorders that in part is due to the fact that *de novo* mutations affect only one copy of each gene; in other words, the functional mutations have dominant phenotypes. Genes sensitive to LoF or missense mutations influence different types of sensitive pathways that are involved in brain development and function. Our study shows that the analysis of functional *de novo* mutations is not only influenced by disorder-specific mechanisms, but also captures the processes that are sensitive to specific class of mutations. Thus, we suggest that previous functional studies of neuropsychiatric disorders may have reached specific conclusions based on which classes of mutations they studied.

Our results indicate that the mechanisms leading to specific disorders are sensitive to different class of mutations. The sensitive processes, shared across disorders occur at different levels. First, at the molecular level, highly connected ‘hub genes’ are more sensitive to perturbations (Jeong et al., 2001). A gene can be highly connected because the protein product has many interactions with other proteins or because it has a role in regulation of multiple genes via transcription or translation. Our analysis suggests that genes involved in neuron development that are also highly connected in the PPI network are more affected by missense mutations, possibly because those mutations affect protein structure or function. Genes sensitive to LoF mutations are characterized by being chromosome and chromatin organizers and by having low variation in gene expression.

In the analysis of gene co-expression, the correlation between ID and ASD depended on mutation class. It suggests that genes intolerant to specific types of functional mutations are co-expressed during development. For instance, the M2 module that is enriched for chromatin regulators was unsurprisingly associated specifically with genes with LoF mutations in both ASD and ID. The three modules that are enriched for genes with functional mutations in ASD are also enriched with constrained genes. Thus, previous findings that genes with LoF mutations in ASD co-expressed in fetal cortex (Parikshak et al., 2013; Willsey et al., 2013) do not necessarily reveal ASD specific mechanisms, but instead expose the type of genes and processes that are sensitive to LoF mutations. This is also supported by the analysis of ID recessive genes that points to a completely different pathway that is sensitive to mutations affecting both copies of the gene.

Second, at the level of the brain, the analysis of ASD and ID genes together with constrained genes suggests that the cortex, especially during early development, is the most vulnerable region to LoF mutations. It was previously suggested that the cortex might be more vulnerable to genetic mutations because of the intense evolution in the recent lineage leading to humans, and the insufficient time to evolve enough buffering capacity (McGrath et al., 2011). We found that across different brain regions, the genes intolerant to mutations are preferentially expressed in early fetal and to a lesser degree during adolescence, suggesting that those two periods of brain development are the most sensitive to genetic insults.

Third, our analysis across disorders suggests that males are more vulnerable to LoF mutations, but not necessarily to missense mutations. The trend of increasing rate of LoF mutations in females is consistent with biological differences between the sexes and with more robust brain development in females (Suliman et al., 2014).

Despite the extensive convergence across disorders discussed above, we can still identify unique patterns of preferential expression in the brain for the different disorders. In ID, the enrichment of genes with functional mutations was restricted to the cortex in both fetal and young adulthood, depending on the type of mutations. Genes with LoF mutations in ASD showed enrichment not only in fetal cortex, but also in additional brain regions – the cerebellum and striatum. The different enrichment patterns seen in ID, suggest that altered function of multiple brain regions account for the range of cognitive, social and restrictive behaviors seen in ASD.

The analysis of genes located in the 108 SCZ loci suggests that common variants associated with SCZ affect synaptic and neuronal processes that are active in the cortex during adolescence and that influence dopaminergic neurons. These results elucidate potential causative mechanisms, which are consistent with SCZ age of onset as well as with the classical theories of cortical dysfunction and the role of dopamine in SCZ (Howes and Kapur, 2009; Winterer and Weinberger, 2004). Not only that antipsychotic drugs increase presynaptic dopamine metabolism and block dopamine D2 receptors, but also brain imaging and postmortem studies have found strong evidence for a dopaminergic hyper-function in the striatum of patients with SCZ (Howes and Kapur, 2009; Winterer and Weinberger, 2004).

Our analysis provides evidence that dopamine is not only part of the pathophysiology of the disorder but also involved in its etiology. Since extensive GWAS loci are not available for ASD or ID, we cannot directly test the specificity of the results. However, a recent GWAS study of educational attainment, which is genetically correlated with cognitive performance (but not with SCZ), found the candidate genes to preferentially expressed in the brain during the prenatal period (Okbay et al., 2016). The preferential expression of SCZ genes in adolescent cortex and educational attainment in prenatal period are consistent with a close relationship between the temporal expression of the genes and the phenotypic manifestation of the genetic variants.

## Methods

### Data Collection

The *de novo* mutations analyzed in this paper were collected from 14 different studies: (1) mutations found in children with ASD and their unaffected siblings (Iossifov et al., 2012, 2014; Neale et al., 2012; O’Roak et al., 2012; De Rubeis et al., 2014; Sanders et al., 2012), (2) mutations found in SCZ patients and their unaffected siblings (Fromer et al., 2014; Girard et al., 2011; Gulsuner et al., 2013; McCarthy et al., 2014; Xu et al., 2008), (3) mutations found in ID patients (Gilissen et al., 2014; Hamdan et al., 2014; de Ligt et al., 2012; Rauch et al., 2012), and (4) mutations found in control families (Rauch et al., 2012; Xu et al., 2008). We annotated all the mutations using Annovar (Wang et al., 2010) with the Ensembl annotation, and included only protein coding mutations for further analysis. In total, we analyzed 4,481 ASD mutations, 309 ID mutations, 975 schizophrenia mutations, 1,931 mutation in unaffected siblings, and 44 mutations in control families. The mutations were grouped into LoF, missense and synonymous mutations. LoF mutations included frameshift, stop codon and splicing mutations. Missense mutations were defined as non-synonymous SNVs and indels, which were not LoF.

### De novo mutations rates

The average number of *de novo* mutations per individual was calculated for each disorder and each mutation category (LoF, missense and synonymous). To control for factors that influence estimates of absolute rates of *de novo* mutations we normalized the number of functional mutations (LoF or missense) to the number of synonymous mutations identified in each study. We then compared the normalized measure between cases and controls. We excluded from the calculation the study of de Ligt et al. (2012), which has was partly overlapping with Gilissen et al. (2014). In ASD, we used data from two studies that included all the samples from previous work (Iossifov et al., 2014; De Rubeis et al., 2014). A pairwise t-test was used to compare between the per-individual mutation rate in the three disorders and the control sample. In order to test for differences in mutation rates between males and females we used a two-way ANOVA to test for interactions between sex and disease status. For each mutation type, *P*-values were corrected for multiple testing using FDR correction.

### Overlap between conditions

To examine the overlap across different class of mutations, we used the *dnenrich* software (Fromer et al., 2014) to test whether genes with mutations in one disorder were significantly enriched among the genes with same type of mutations in the other disorders. The *dnenrich* permutation framework generates random set of mutations but controls for gene size, structure and local trinucleotide mutation rate. The overlap of mutated genes between the different disorders was tested using 10,000 permutations. For each mutation category, we tested the overlap of the list of genes in each condition with all other conditions. Genes that were mutated more than once in a specific list were given a weight based on the number of occurrences. The program preforms permutations on the entire exome using the original mutation list and creates multiple permutated gene lists, controlling for GC content and gene length. The degree of overlap was calculated by dividing the observed number of overlapping genes by the number of overlaps in the permutations. For each mutation type, *P*-values were corrected for multiple testing using FDR correction.

### Analysis of constrained genes

We used levels of gene constraint, and the list of genes with significant constraint from Samocha et al. (Samocha et al., 2014). We calculated the mean constraint level for genes with mutations in each disorder using the constraint score available for missense mutations (presented as z scores). ANOVA was used to test for differences in constraint score, followed by pairwise t-test with FDR correction (*pairwise.t.test* function in R). We tested the enrichment of the 1,003 constrained genes among the genes mutated in each disorder relative to the control using Fisher’s exact test with FDR correction.

### Protein–protein interactions

Protien-protein interaction (PPI) data was downloaded from the Mentha project (Calderone et al., 2013) (on August 24, 2015). The dataset contains an integrated PPI information with a reliability score for each interaction. We used the reliability score to generate a weighted measure of the number of interactors of each protein by summing the reliability score of each of the protein interactions. We then tested the difference between the weighted average number of interactions for genes mutated in the different disorders relative the control, and across the different mutation categories. Significance was based on a Kolmogorov–Smirnov test, followed by FDR correction. Similarly, we tested for the difference between the 1,003 constrained genes and the unconstrained genes.

### Variation in gene expression

We used a previously published single-cell gene expression from human brain (Darmanis et al., 2015). The data was comprised of 465 samples. We converted the raw read counts to counts per million (cpm) and normalized to gene exonic length. We divided the dataset to different cell types (Astrocytes, Endothelial, Microglia, Neuron, Oligodendrocytes and Oligodendrocyte progenitors) by clustering them using cell type-specific marker genes, which were used in the original study (Darmanis et al., 2015). We calculated a standardized measure of expression variation controlled for the mean expression based on the coefficient of variation (CV) (standard deviation divided by mean) for genes expressing (cpm > 0) in more than 10% of the 247 neuronal cells. Since the log of the CV showed linear correlation with the log of the average expression, we calculated the residuals from the linear regression as a standardized measure of expression variation. ANOVA was used to test for the differences between the standardized measures across conditions. *P*-values were adjusted by FDR correction.

### Gene expression enrichment in specific brain regions

We used a previously published method (Dougherty et al., 2010; Xu et al., 2014), which is based on a specificity index probability (pSI) statistic – a measure of enrichment of a gene in a tissue or cell type. We used the pSI R package to calculate the enrichment *P*-values for gene lists in different brain regions (Amygdala, Cerebellum, Cortex, Hippocampus, Striatum, Thalamus), and across different developmental stages (early fetal, early mid fetal, late mid fetal, late fetal, neonatal-early Infancy, late Infancy, early childhood, middle-late childhood, adolescence, young adulthood). Significance of the overlap with the list of genes with *de novo* mutations was calculated using the *dnenrich* program (Fromer et al., 2014) with 10,000 permutations. An FDR procedure was used to correct for multiple regions and stages.

### Gene co-expression analysis in the cortex

We used an approach that was previously applied to identify convergence in ASD-associated genes (Ben-David and Shifman, 2012; Parikshak et al., 2013) based on weighted gene co-expression network analysis (WGCNA) (Zhang and Horvath, 2005). The approach relies on the assumption that co-expressed genes are functionally related. By dividing the genes in the genome into modules of co-expressed genes, it is possible to test the enrichment of risk genes in each module (Ben-David and Shifman, 2012; Parikshak et al., 2013). We used a published WGCNA that was based on gene expression from the developing human cortex (PCW 8 – 12 months) (Parikshak et al., 2013). The network was comprised of 22,084 genes mapped to 18 modules. The enrichment analysis was performed only on protein coding genes (n = 15,591). For each mutation class and each disorder, we tested the enrichment in each module using *dnenrich* (Fromer et al., 2014) with 10,000 permutations. For the constrained genes, we tested the enrichment using Fisher’s exact test (using fisher.test function in R). *P*-values were corrected for multiple testing using FDR correction across all modules. In order to study the relationship between the disorders and mutation categories we calculated the Pearson correlation between the −log_10_ of the nominal *P*-values (-logP), which was used for hierarchical clustering (using *heatmap.2* function in R). The −logP was used to measure the correlation in enrichment across modules since the effect sizes were highly influenced by the size of the modules. The significance of the difference between correlation coefficients was calculated using Fisher r-to-z transformation.

### Analysis of SCZ GWAS

We used a multistep procedure to prioritize the most likely causal genes. First, we selected a single SNP in each of the 108 loci with the most significant association signal as the most likely causal SNP. The list of the 108 most significant SNPs was extracted from Ripke et al. (Ripke et al., 2014). Second, assuming that most regulatory variants are within a relatively short distance from the gene we included only genes overlapping 100 kb window around the most significant SNP. RefSeq genes overlapping the 100kb windows around the SNPs were downloaded from the USCS genome browser. On average, there were around two genes per window, totaling 210 RefSeq genes (including non-coding RNAs) (Table S4). Third, we prioritize the genes based on the assumption that SCZ risk genes share functional annotations and will be more functionally similar to each other relative to other genes in the list. The list of genes were tested for annotation enrichment using the ToppGene Suite (Chen et al., 2009). Genes that were included in at least one of the significant annotations were considered as the most likely SCZ candidate genes. One of the GWAS loci includes three genes that are a part of the nicotinic receptor cluster (CHRNA5-CHRNA3-CHRNB4). To correct for possible bias due to duplicated genes that are physically located within the same loci we treated them as a single entity. Specifically, to avoid the overrepresentation of nicotinic receptor genes we included only one representative gene in the analysis. Control gene lists were created by shifting the 100 kb windows by one Mb or by selecting random sets of 108 SNPs that were found to be significantly associated with other unrelated traits in GWASs. No significant enrichment was found for simulated control gene lists. Tests for enrichment of genes in brain regions or cell types was as described above using previously published method and data (Dougherty et al., 2010).

## Acknowledgments

We thank Jonathan Flint for his valuable comments on the manuscript. This research was supported by the National Institute for Psychobiology in Israel – founded by The Charles E. Smith Family and by the Israel Science Foundation (grant no. 688/12). Eyal Ben-David was supported by the Dennis Weatherstone Pre-doctoral Fellowship from Autism Speaks (grant no. 8595).

## Supplemental Figures and Figure legends

Figure S1, related to Figure 1

**Figure S1 (related to Figure 1).**
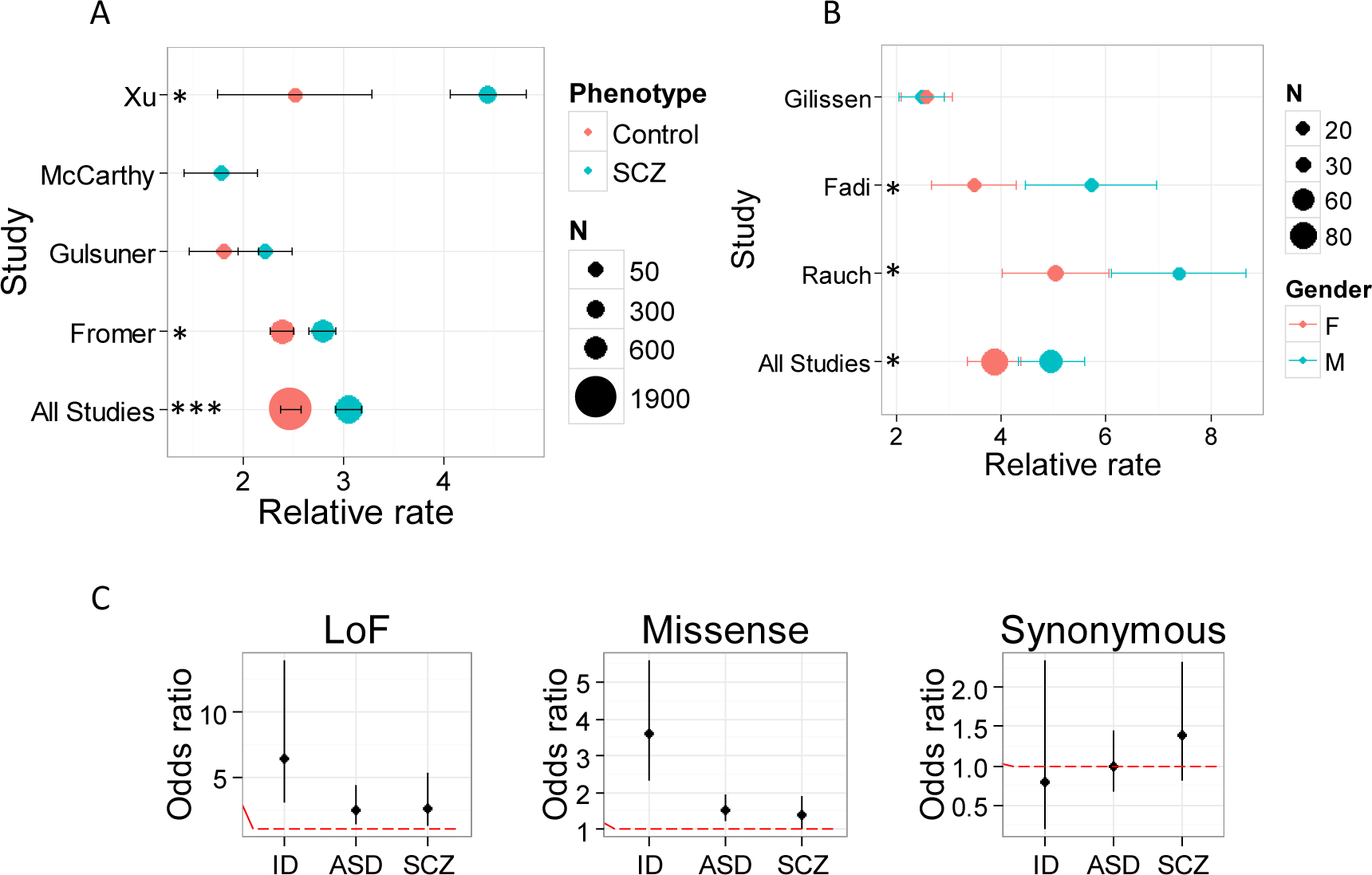
Rates of *de novo* missense mutation per study and overlap with genes under selective constraint. (A) Rates of missense mutation in SCZ and matching controls. Values are mean rate of missense mutations relative to synonymous mutations (± SEM). The size of the symbols is proportional to the sample size. Test of significance are for differences in missense mutation rate between SCZ and controls used in each study (Fromer et al., 2014; Gulsuner et al., 2013; McCarthy et al., 2014; Xu et al., 2008). *, *P* < 0.05. ***, *P* < 0.001. (B) Rates of missense mutations in males and females with ID across studies (Gilissen et al., 2014; Hamdan et al., 2014; Rauch et al., 2012). Values are mean rate of missense mutations relative to synonymous mutations (± SEM). The size of the symbols is proportional to the sample size. *, Significant interaction between sex and disease status (*P* < 0.05). (C) Association between 1003 constrained genes and genes with *de novo* mutations. Values are odds ratio relative to the control (± 95% confidence interval). The red dotted line indicates an odds ratio of 1.

Figure S2, related to Figure 2

**Figure S2 (related to Figure 2).**
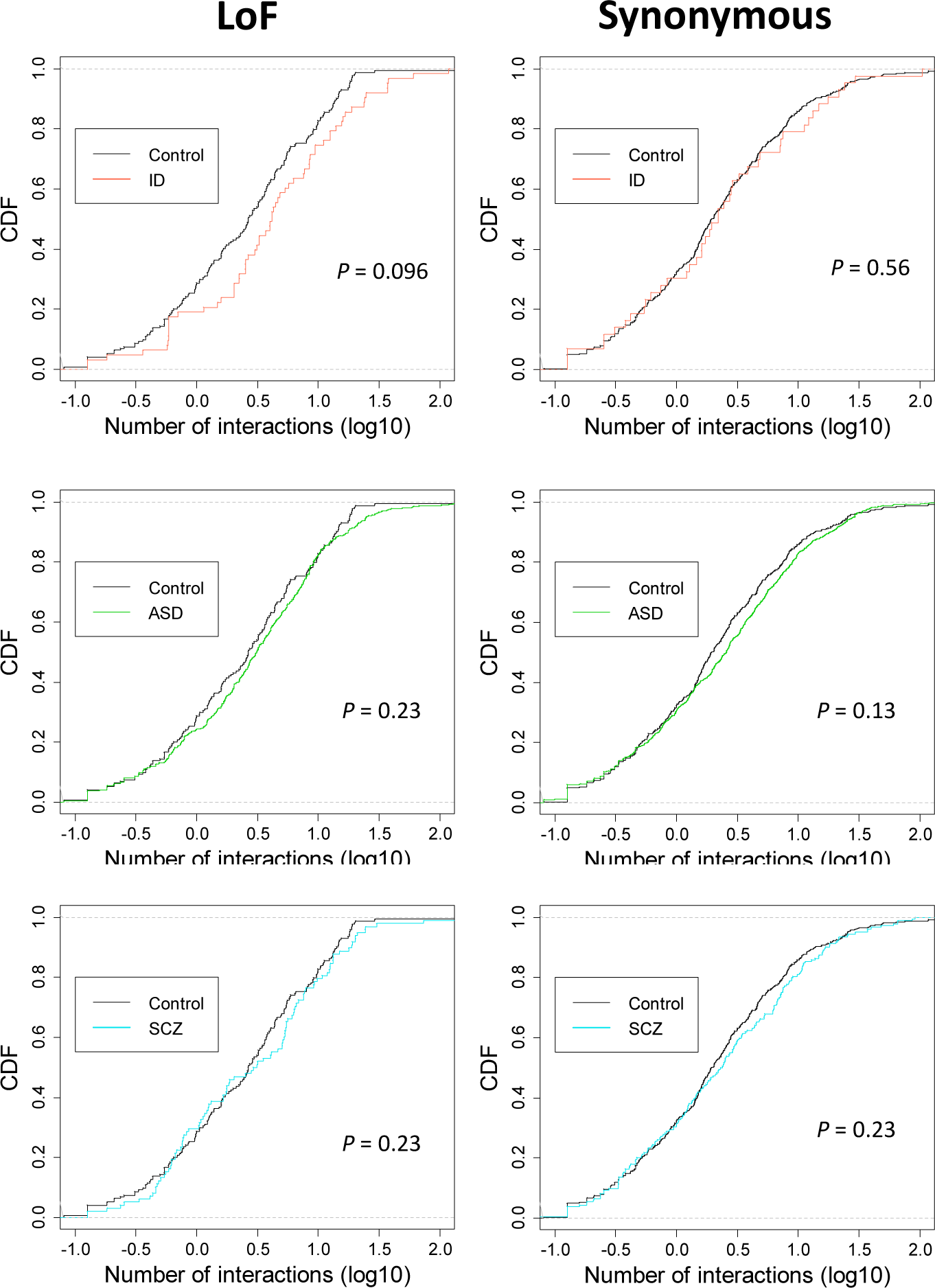
Number of protein-protein interactions for genes with LoF and synonymous mutations. Cumulative distribution function (CDF) of the number of interactions (log_10_) plotted for genes with LoF mutations (left) and synonymous mutations (right) in ID, ASD and SCZ relative to control. *P*-values were calculated using a two-sample Kolmogorov–Smirnov test, and corrected by FDR.

Figure S3, related to Figure 3

**Figure S3 (related to Figure 3).**
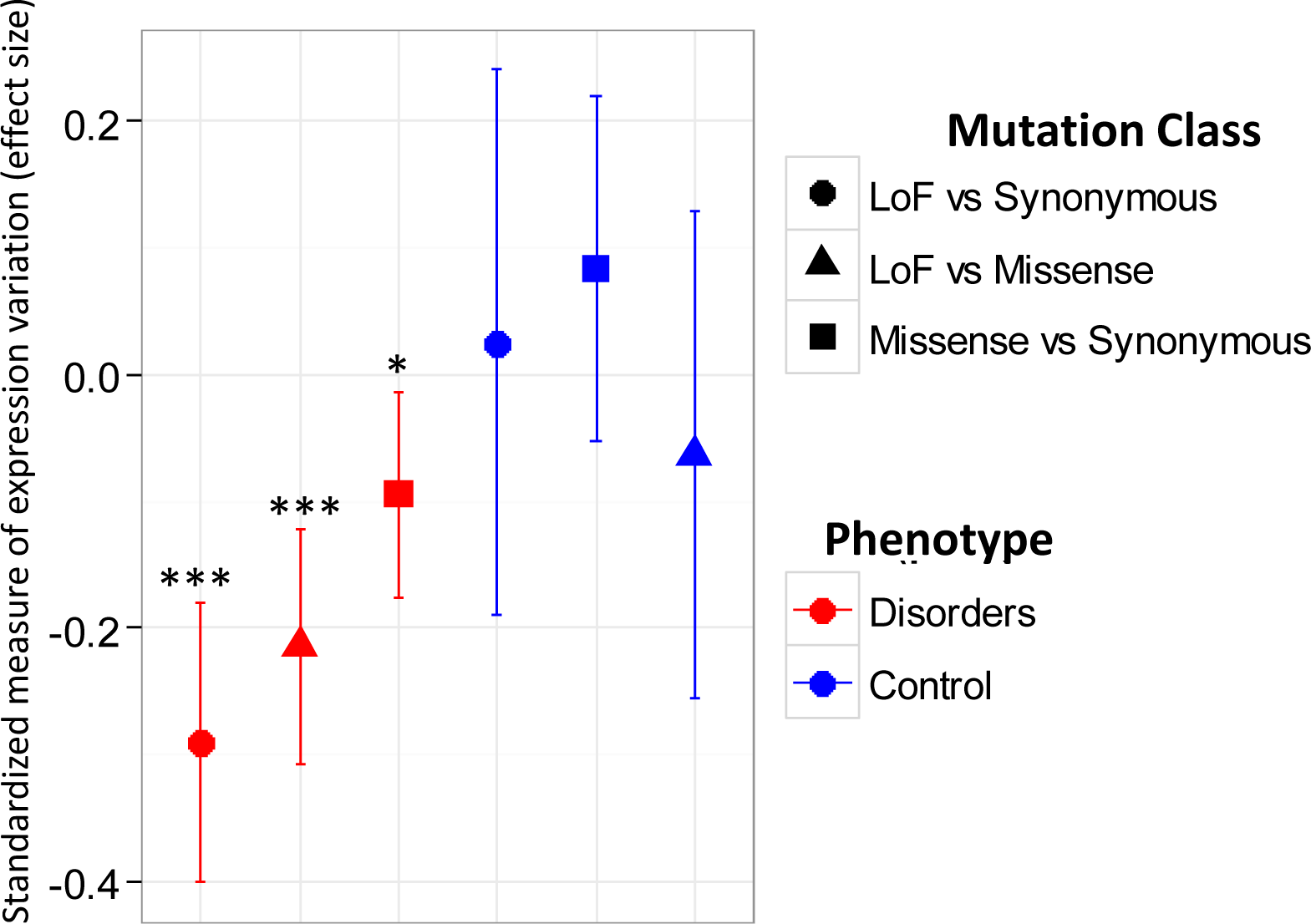
Genes harboring functional mutations in the disorders show less vitiation in gene expression, especially genes with LoF mutations. Values are the effect size (Cohen’s d) of the difference in expression variation between different mutation classes. Error bars are 95% confidence interval. *, *P* < 0.05. ***, *P* < 0.001.

Figure S4, related to Figure 5

**Figure S4 (related to Figure 5).**
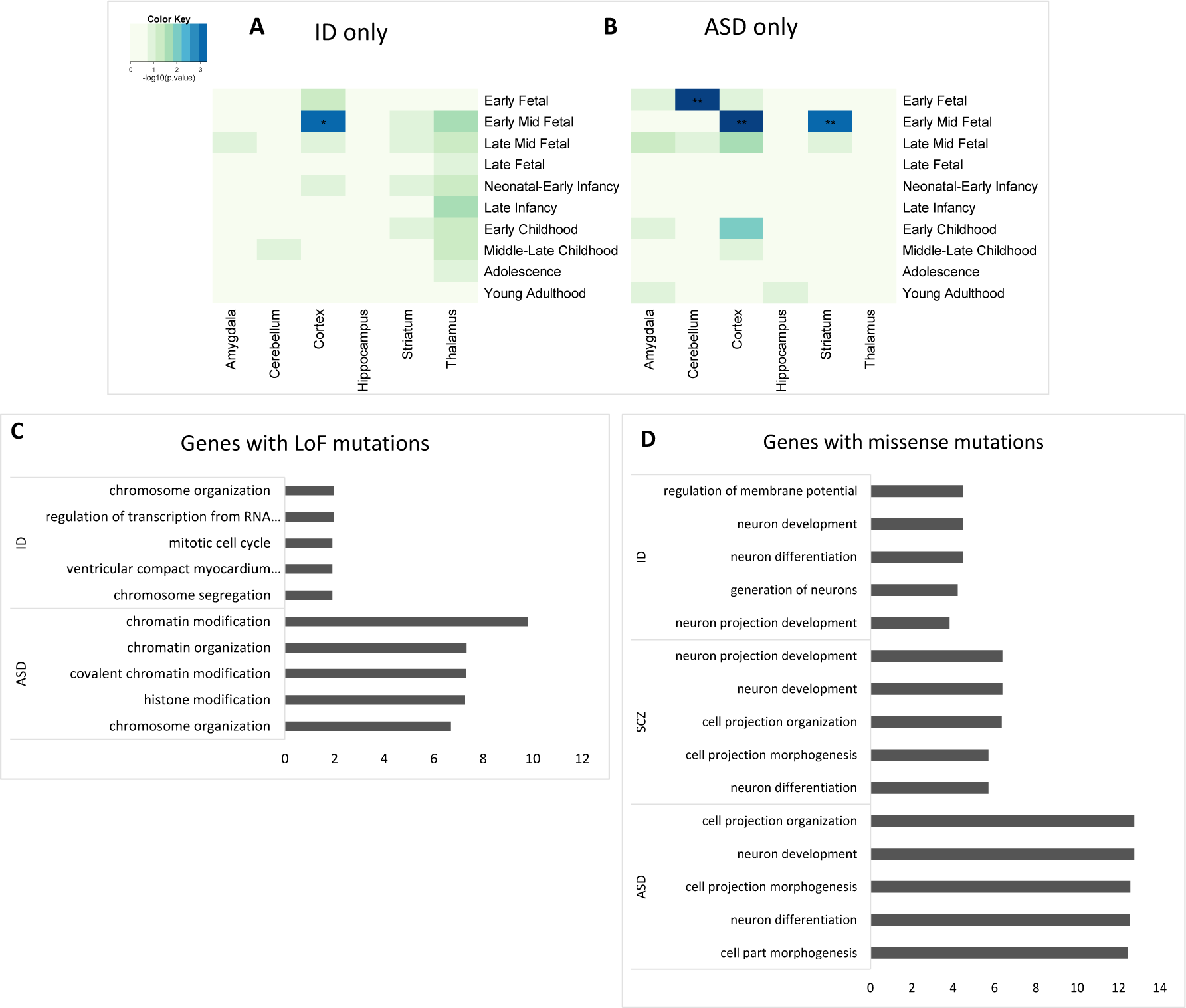
Region and stage specific enrichment in the human brain and functional enrichment analysis of genes with *de novo* mutations. (A) Enrichment analysis of genes with LoF mutations in different regions of the human brain and during different stages of development for genes with mutations only in ID (non-overlapping genes) or (B) ASD-specific genes. Heatmap colors represent the enrichment significance presented as −log_10_ of the *P*-values. *P*-values are calculated based on permutations in the *dnenrich* software and are corrected for multiple tests using FDR. *, FDR corrected P < 0.05. **, FDR corrected P < 0.01. (C) Functional enrichment analysis of genes with LoF mutations, and (D) missense mutations. Top five significantly enriched biological processes are shown for each disorder. The analysis was based on the pathway enrichment tool, ToppGene Suite. Values are −log_10_ of the FDR corrected *P*-values.

Figure S5, related to Figure 6

**Figure S5 (related to Figure 6).**
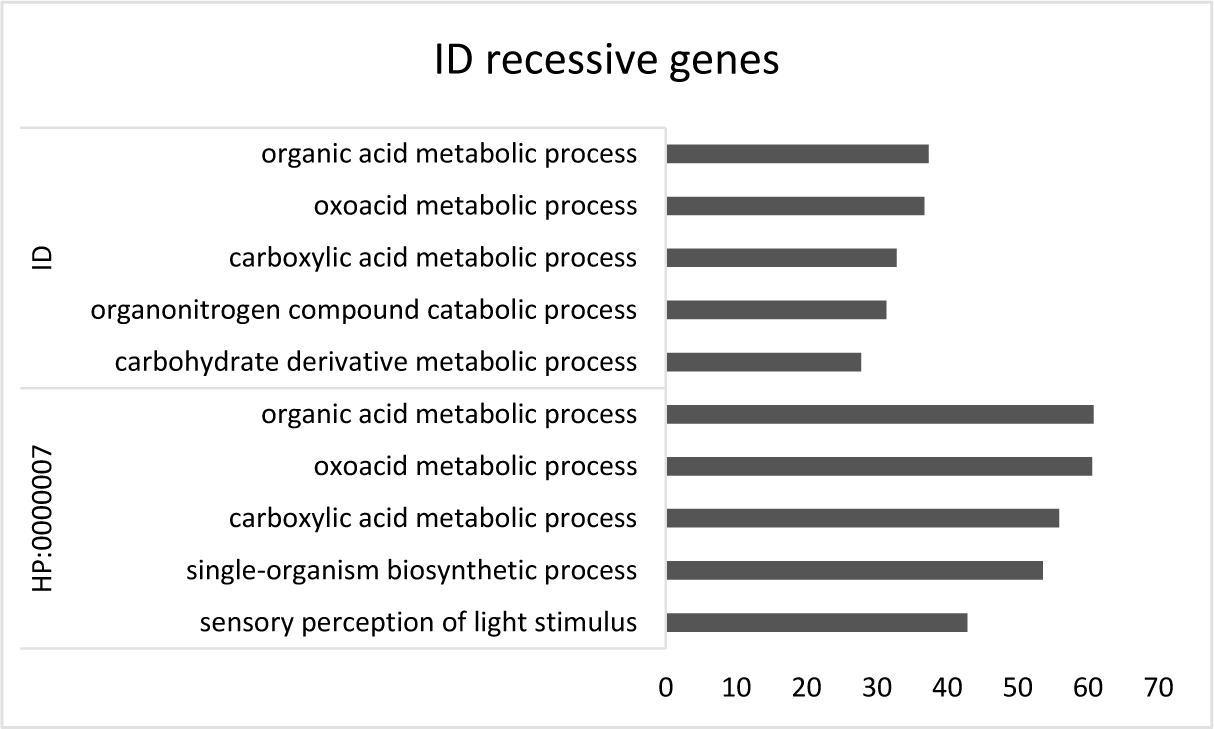
Functional enrichment analysis of ID genes with autosomal recessive inheritance (top) and all genes with autosomal recessive inheritance (HP:0000007) based on the Human Phenome Ontology (bottom). Top five significantly enriched biological processes are shown for each. The analysis was based on the pathway enrichment tool, ToppGene Suite. Values are −log_10_ of the FDR corrected *P*-values.

Figure S6, related to Figure 7

**Figure S6 (related to Figure 7).**
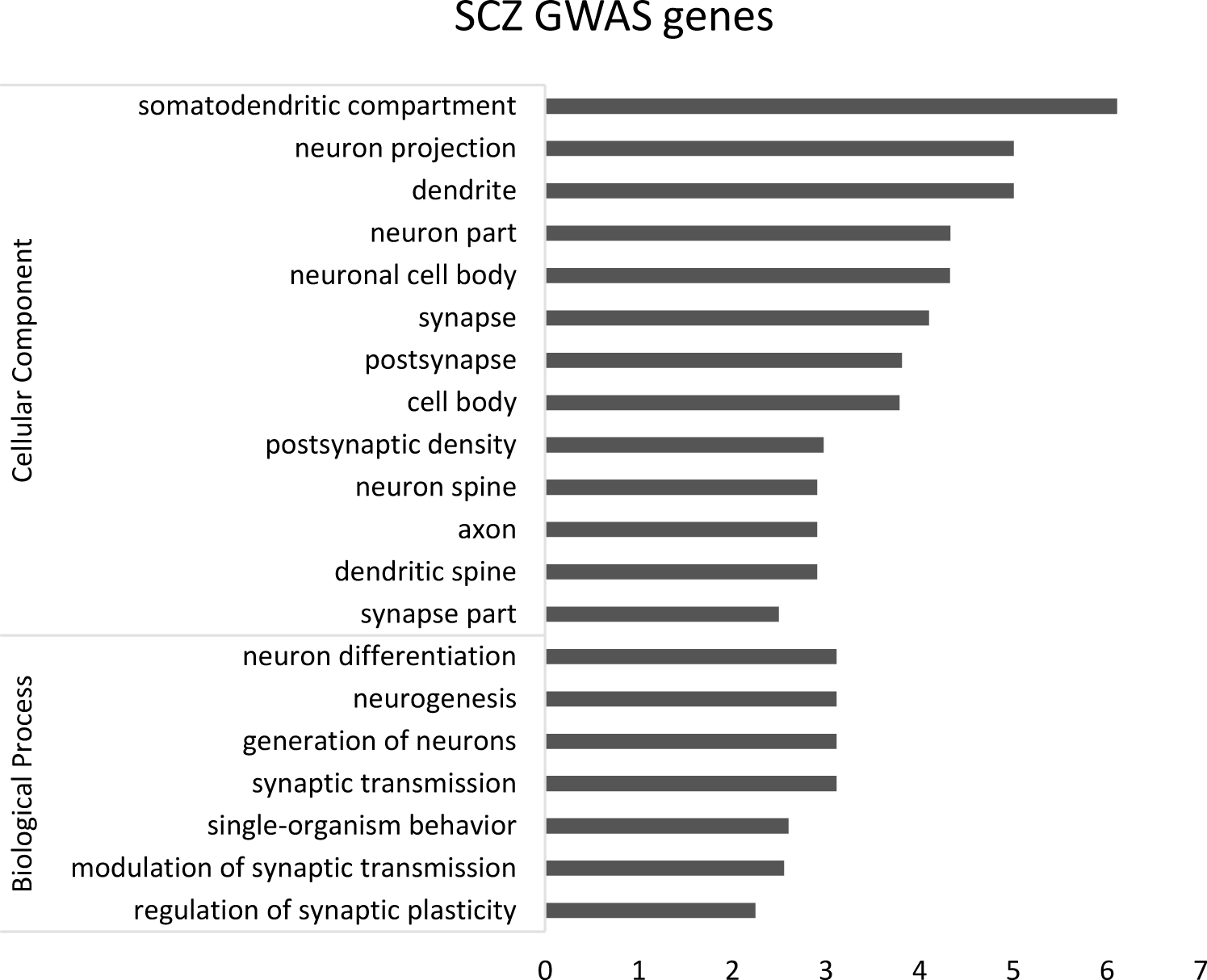
Functional enrichment analysis of genes implicated in schizophrenia GWAS. Significantly enriched cellular component and biological processes for genes in 100 kb window around the most significant SNPs in schizophrenia GWAS. The analysis was based on the pathway enrichment tool, ToppGene Suite. Values are −log_10_ of the FDR corrected *P*-values.

## Supplemental Tables

**Table S2.** Overlap between genes with *de novo* mutations across conditions

| Conditions |  | LoF |  | Missense |  | Synonymous |  |
| --- | --- | --- | --- | --- | --- | --- | --- |
|  |  | O/E | Corrected *P* | O/E | Corrected *P* | O/E | Corrected *P* |
| ID | ASD | 15.24 | 0.00030 | 2.24 | 0.00017 | 0.88 | 0.84 |
| ID | SCZ | 10.06 | 0.00063 | 2.71 | 0.00017 | 0.86 | 0.84 |
| ID | Control | 0.00 | 1.0 | 1.56 | 0.0011 | 0.37 | 0.93 |
| ASD | SCZ | 2.94 | 0.0013 | 2.80 | 0.00017 | 1.19 | 0.71 |
| ASD | Control | 1.80 | 0.040 | 1.43 | 0.016 | 1.15 | 0.71 |
| SCZ | Control | 0.91 | 0.81 | 1.05 | 0.47 | 1.12 | 0.81 |

O/E, observed to expected ratio. Corrected P, P values generated by permutations with the *dnenrich* software (Fromer et al., 2014) and corrected with the Benjamini and Hochberg false discovery rate (FDR) procedure.

**Table S3.** Differences in mean constraint score between conditions (related to Figure 1C)

| Mutation type | ANOVA *P* value | Post-hoc pairwise t test with FDR correction |  |  |  |  |  |
| --- | --- | --- | --- | --- | --- | --- | --- |
|  |  | ASD-Control | ASD-ID | ASD-SCZ | ID-Control | ID-SCZ | SCZ-Control |
| LoF | 4.9E-09 | 1.3E-05 | 3.0E-04 | 0.25 | 8.2E-09 | 2.4E-04 | 0.035 |
| Missense | 6.7E-08 | 5.0E-04 | 5.2E-05 | 0.82 | 7.8E-08 | 1.2E-04 | 0.025 |
| Synonymous | 0.39 | - | - | - | - | - | - |

